# A conserved role for stomatin domain genes in olfactory behavior

**DOI:** 10.1101/2022.09.26.508537

**Authors:** Xiaoyu Liang, Morgan Taylor, Rebekah Napier-Jameson, Canyon Calovich-Benne, Adam Norris

## Abstract

The highly-conserved stomatin domain has been identified in genes throughout all classes of life. In animals, different stomatin domain-encoding genes have been implicated in the function of the kidney, red blood cells, and specific neuron types, though the underlying mechanisms remain unresolved. In one well-studied example of stomatin domain gene function, the *C. elegans* gene *mec-2* and its mouse homologue *Stoml3* are required for the function of mechanosensory neurons, where they modulate the activity of mechanosensory ion channels on the plasma membrane. Here we identify an additional shared function for *mec-2* and *Stoml3* in a very different sensory context, that of olfaction. In worms, we find that a subset of stomatin domain genes are expressed in olfactory neurons, but only *mec-2* is strongly required for olfactory behavior. *mec-2* acts cell-autonomously and does not have any observable effect on olfactory neuron development or morphology, but modestly reduces olfactory neuron activity. We generate a *Stoml3* knockout mouse and demonstrate that, like its worm homologue *mec-2*, it is required for olfactory behavior. In mice, *Stoml3* is not required for odor detection, but is required for odor discrimination. Therefore, in addition to their shared roles in mechanosensory behavior, *mec-2* and *Stoml3* also have a shared role in olfactory behavior.

## INTRODUCTION

Olfaction is a remarkable sensory system, enabling animals to detect and distinguish among a very large range of odors, and at very low concentrations. For example, it was recently estimated that humans are able to distinguish as many as a trillion distinct odors^1^, in some cases at thresholds below 1 part per billion^2^. A number of cellular and molecular aspects of the olfactory system of various animals have been well characterized, including odorant-receptor interactions, signal transduction cascades within olfactory neurons, and downstream neuronal circuits^3–6^. However, many factors that are highly expressed in olfactory neurons remain completely uncharacterized or with poorly defined function^7–9^.

We recently found that the stomatin domain gene *mec-2* is highly expressed in *C. elegans* olfactory neurons, and is required for olfaction^10^. The canonical role for *mec-2* and its mouse homologue *Stoml3* is as a component of the mechanotransduction machinery expressed in mechanosensory neurons^11,12^. Our recent results therefore suggest that *mec-2*, and perhaps *Stoml3* or other stomatin domain genes, might have previously unappreciated roles in olfactory neurons.

*mec-2* and *Stoml3* belong to a family of proteins defined by the presence of a highly-conserved stomatin domain, which can be found across all domains of life^13^. The typical architecture of stomatin domain genes entails a central conserved stomatin domain and a hydrophobic region that mediates membrane association (Fig 1A). These are flanked by divergent N- and C-termini, which can confer specific functional and regulatory features^10,14,15^. Stomatin domain proteins localize to the plasma membrane via their hydrophobic region, where some family members have been shown to modulate the activity of membrane proteins, though the precise mechanisms by which they do so remain elusive^14,16,17^.

**Figure 1:**
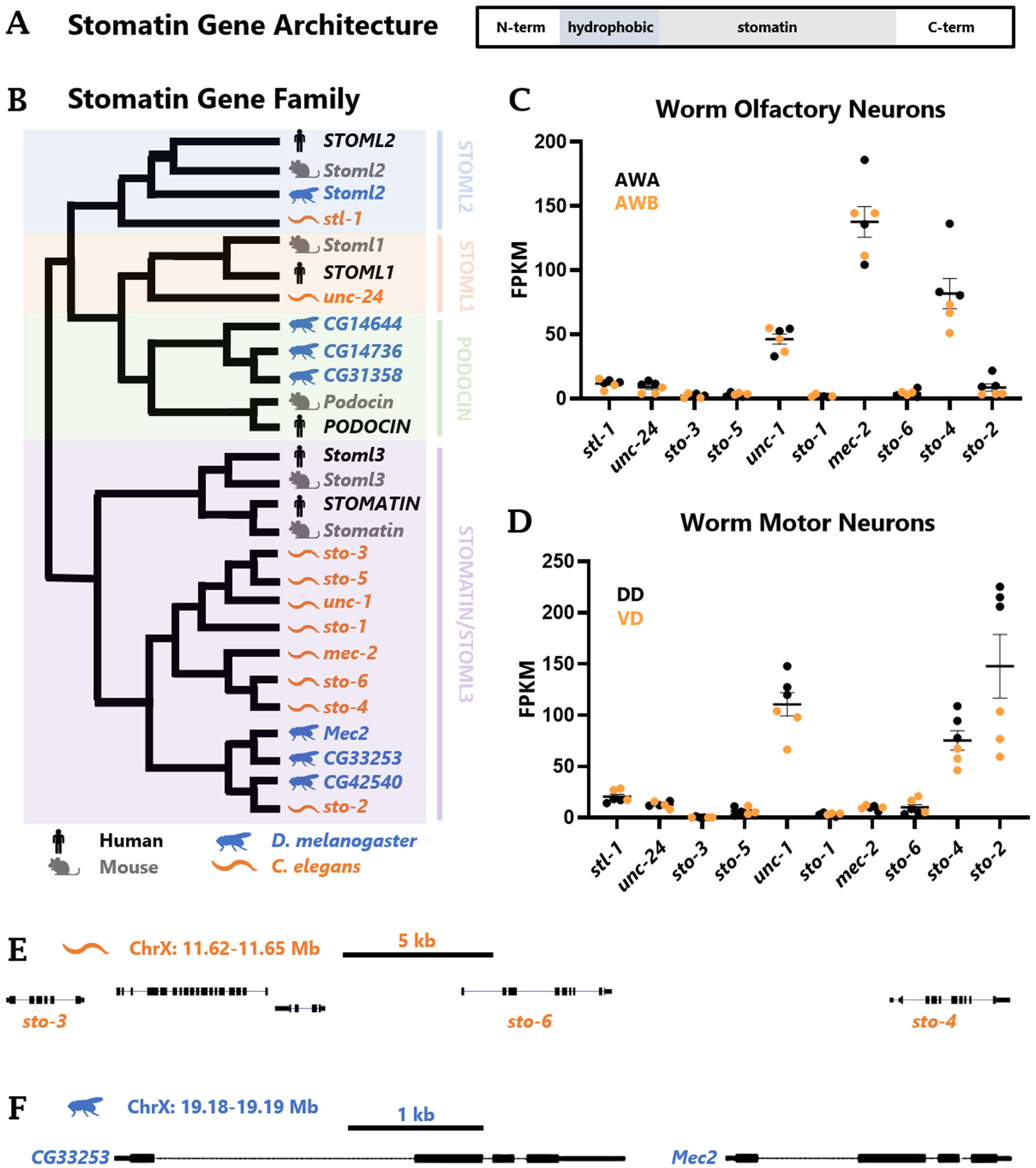
A subset of stomatin domain genes are expressed in *C. elegans* olfactory neurons. (A) Domain organization of stomatin domain genes. Colored in shades of gray are the central conserved stomatin domain and the hydrophobic domain that mediates association with the membrane. Flanking these are the divergent N- and C-termini (B) Gene tree for family of stomatin domain gene homologues, TF105750 (stomatin-like 2) from TreeFam^20^. Some organism silhouettes created with BioRender.com. (C-D) RNA Seq data from CenGEN consortium sorted neuron populations, mapped to the worm genome with STAR and gene reads counted by HTSeq. (C) Olfactory neurons AWA and AWB, and (D) Motor neurons DD and VD. (E) Cluster of three *Stoml3/Stomatin-like* genes in the *C. elegans* genome on chromosome X. (F) Two immediately adjacent *Stoml3/Stomatin-like* genes in the *Drosophila* genome on chromosome X.

Here we show that both worms and mice express a small subset of stomatin domain genes in olfactory neurons, and that a single stomatin domain gene, *mec-2*, is strongly required for olfactory behavior in worms. We demonstrate that *mec-2* acts in a cell-autonomous manner, and that multiple alternatively spliced *mec-2* isoforms can substitute for each other. We also generate a *Stoml3* knockout mouse and demonstrate that, like its homologue *mec-2* in worms, *Stoml3* is required for proper olfactory behavior in mice. Specifically, *Stoml3* KO mice are able to detect odors, but are unable to efficiently distinguish between odors. Therefore, in addition to their conserved roles in mechanosensory behavior, *mec-2* and *Stoml3* have an additional conserved role in olfactory behavior in both worms and mice.

## RESULTS

### Conserved and species-specific stomatin domain genes

We recently showed that the *C. elegans* stomatin domain protein MEC-2 is expressed in olfactory neurons in addition to its previously-described expression in mechanosensory neurons^10^. We also showed that *mec-2* mutants are deficient in both mechanosensory and olfactory behaviors^10^. The worm genome contains nine additional stomatin domain genes (Fig 1B), and while *mec-2* has been extensively characterized in *C. elegans*, many of the other stomatin domain genes remain uncharacterized. To test whether additional stomatin domain genes are involved in olfaction, we first determined phylogenetic relationships among stomatin domain proteins in *C. elegans* and other well-studied metazoan genomes (Fig 1B), with the aim of determining whether closely related genes are co-expressed in specific neuron types (Fig 1C-D).

We noticed a striking expansion of specific stomatin domain gene members in both *Drosophila* and *C. elegans*. For example, a single *Stoml2* orthologue exists in humans, mice, flies, and worms. On the other hand, the *Stomatin/Stoml3* sub-family contains two genes in humans and mice, three in flies, and eight in worms (Fig 1B). This observation led us to examine whether there is evidence for recent expansion of stomatin domain genes in *Drosophila* or *C. elegans*. Indeed, three of the eight *C. elegans Stomatin/Stoml3* homologues reside in a cluster on chromosome X (Fig 1E). Such clustering is a hallmark of recently-duplicated genes in *C. elegans*^18,19^. The remaining *C. elegans Stomatin/Stoml3* homologues do not reside in genomic clusters, but are all located on the X chromosome, while the more distantly-related Stomatin domain genes are distributed across autosomes.

*Drosophila* stomatin domain genes also exhibit genomic clustering: two of the three *Stomatin/Stoml3* homologues are located immediately next to each other on chromosome X (Fig 1F), as are two of the three closely-related *Drosophila* homologues of *Podocin* on chromosome 3 (Fig S1). Together these results indicate that *C. elegans* and *Drosophila* genomes contain expanded repertoires of stomatin domain genes, most notably in genes related to *Stomatin*/*Stoml3*.

### Subset of stomatin domain genes expressed in *C. elegans* olfactory neurons

We analyzed RNA Seq data generated from specific FACS-sorted neuron types^21^ to determine which of the *C. elegans* stomatin domain genes are expressed in olfactory neurons. We focused on the AWA and AWB neurons, which are involved in detecting attractive and repulsive odors, respectively^22^. We previously showed that *mec-2* is required for detection of both AWA- and AWB-specific odorants^10^. Cell-specific RNA Seq confirms that *mec-2* is highly expressed in both AWA and AWB neurons (Fig 1C). As a comparison, we also assayed expression of stomatin domain genes in an additional class of neurons, the VD and DD motor neurons. In contrast to olfactory neurons, *mec-2* expression is nearly undetectable in motor neurons (Fig 1D). This demonstrates that *mec-2* is expressed in specific subsets of neurons, including the AWA and AWB olfactory neurons.

We also detected substantial levels of expression for both *sto-4* and *unc-1* in AWA and AWB olfactory neurons (Fig 1C). In contrast to *mec-2*, however, *sto-4* and *unc-1* are also highly expressed in motor neurons, suggestive of a broader function in various neuron types (Fig 1D). Indeed, *unc-1* was previously shown to be expressed and active in a variety of muscles and neurons, including motor neurons^23^. Finally, we note that within the functional classes of neurons we tested, stomatin domain genes tend to exhibit similar expression levels between individual cell sub-types (*e.g. mec-2* is highly expressed in both AWA and AWB). The one exception is *sto-2*, which is highly expressed in motor neurons, but with consistent >3-fold higher expression in DD than in VD motor neurons (Fig 1D). Together these data reveal a diversity of stomatin domain genes in the *C. elegans* genome, which exhibit cell-type specific gene expression within the nervous system.

### Specific stomatin domain genes required for *C. elegans* olfactory behavior

We next tested whether the Stomatin domain genes expressed in olfactory neurons are required for olfactory behavior. Using standard chemotaxis assays (Fig 2A) we previously showed that *mec-2* is required for chemotaxis responses to a variety of volatile odorants, both attractive and repulsive^10^ (Fig 2B-C). We obtained multiple loss-of-function alleles for both of the olfactory-neuron-expressed stomatin domain genes, *unc-1* and *sto-4*. We found that mutations in both genes cause reduced locomotion, which complicates the interpretation of standard chemotaxis assays, which require coordinated locomotion over time toward or away from the olfactants. We therefore altered the duration of the chemotaxis assay from the traditional 1 hour to an extended 2 hours to give these mutants more time to chemotax. Even so, both *unc-1* and *sto-4* mutants exhibit chemotaxis defects (Fig 2B-C), though we reasoned that this still might be largely attributable to locomotory deficits.

**Figure 2:**
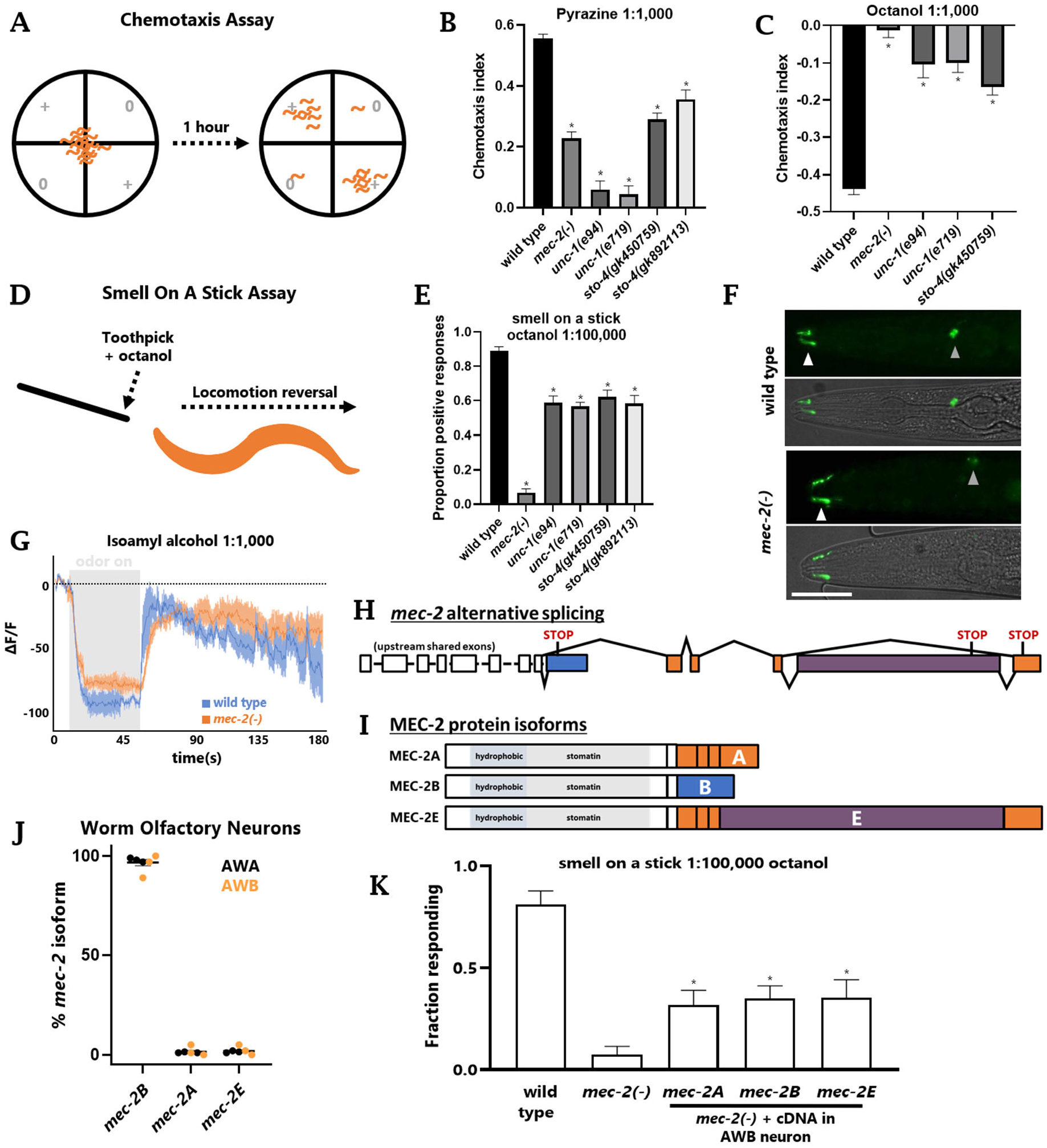
Specific stomatin domain genes required for olfaction in *C. elegans*. (A) Schematic of the chemotaxis assay. Worms are placed in the center of an unseeded worm plate, given a choice between point sources of odor (represented by +) and unscented diluent (represented by 0), and given one hour to chemotax. (B-C) Chemotaxis assays for the attractive odor pyrazine (1:1,000) (B) or the repulsive odor octanol (1:1,000) (C). The chemotaxis index = ((worms in + quadrant)-(worms in 0 quadrant))/(worms in all quadrants), with a maximum value of 1 (attractive) and minimum value of −1 (repulsive). (D) Schematic of the smell on a stick assay for repulsive odors. (E) Smell on a stick assay for octanol. (F) AWA neurons labeled by translational ODR-10::GFP fusion. White arrowheads denote sensory cilia (where ODR-10 localizes). Gray arrowheads denote cell bodies. The cell body in the *mec-2* image is slightly out of the plane of focus. Scale bar represents 50 μm. (G) GCaMP5 imaging of AWC(on) neurons using a microfluidic “olfactory chip.” Isoamyl alcohol (1:1000) administration represented by the gray box. (F) Schematic of *mec-2* alternative splicing and (G) subsequent protein isoforms. (H-I) *mec-2* alternative isoforms at the level of pre-mRNA splicing (H) and protein isoforms (I). The isoforms affect the divergent C-terminus of *mec-2*. (J) CenGEN RNA Seq data reveals *mec-2* is found primarily as the *mec-2B* isoform in AWA and AWB olfactory neurons. (K) Smell on a stick assay, 1:100,000 octanol, statistical test (ANOVA) performed against *mec-2(-)* mutants. Re-expression of any *mec-2* isoform in AWB neuron partially rescues response. Asterisks indicate p < 0.05, one-way ANOVA.

To disentangle the effects of *unc-1* and *sto-4* on olfaction versus locomotion, we turned to a “smell on a stick” assay, in which individual worms are directly presented with an aversive odor, causing wild-type worms to quickly perform a locomotion reversal to avoid the odor (Fig 2D). This assay requires much less coordination of locomotion, allowing us to focus on the olfactory abilities of the mutants. *mec-2* mutants exhibit strong defects in this assay when presented with the repellant odor octanol, while *sto-4* and *unc-1* mutations cause only minor olfactory defects (Fig 2E). From these results, we conclude that *mec-2* is the primary stomatin domain gene required for olfaction in *C. elegans*. While *sto-4* and *unc-1* are expressed in olfactory neurons, their requirement for chemotaxis stems largely from locomotory defects. This is consistent with their high levels of expression in motor neurons as well (Fig 1C).

In mechanosensation, *mec-2* has been shown to affect activity of mechanosensory channels, but not the development or structure of the mechanosensory neurons themselves^24^. We likewise wanted to test whether *mec-2* plays a role in the development or activity of olfactory neurons. We examined the structure of the AWA neuron and found no differences between wild type and *mec-2* mutants: their cell bodies, dendrites, and cilia are all positioned appropriately, as is the olfactory receptor ODR-10 (Fig 2F). We therefore conclude that *mec-2* is not required for the development or morphology of the AWA olfactory neuron.

We also tested the activity of olfactory neurons in wild type and *mec-2* animals, using the well-characterized *str-2::GCaMP5* reporter of calcium activity in AWC(on) olfactory neurons^25,26^. As previously described, in wild-type neurons odor presentation results in a robust decrease in calcium, and odor removal results in a robust increase in calcium^25,26^ (Fig 2G). In *mec-2* mutants, the calcium response is broadly similar, but the magnitude of the calcium response is decreased (Fig 2G). The magnitude of the decrease is modest, but statistically significant (Fig S2). From these experiments we conclude that *mec-2* has a modest effect on olfactory neuron activity in response to olfactory stimuli.

Finally, to test whether *mec-2* acts cell autonomously in olfactory neurons, we performed the smell on a stick assay in *mec-2* mutants with cell-specific *mec-2* re-expression. We used the *str-1* promoter to re-introduce *mec-2* to the AWB neuron, which is involved in sensing the aversive odor octanol^27^. We previously demonstrated that *mec-2* encodes three different isoforms with distinct functional properties in different cell types (Fig 2H-I). In agreement with our previous work using reporters^10^, cell-specific RNA Seq indicates that the *mec-2B* isoform predominates in AWA/AWB olfactory neurons (Fig 2J). We generated cDNAs for each of the three isoforms and tested whether each could rescue the olfactory response to octanol when artificially expressed in AWB. Indeed, transgenic re-introduction of any of the three *mec-2* isoforms in the AWB neuron partially rescues octanol avoidance responses in *mec-2* mutants (Fig 2K). This is in contrast with our previous work in mechanosesnroy neurons, which showed that *mec-2B* is not functional, while *mec-2A* and *mec-2E* must be co-expressed for proper mechanosensory behavior^10^. Together, these experiments indicate that all three *mec-2* isoforms are functional when ectopically expressed in AWB, and that *mec-2* acts cell-autonomously in the AWB olfactory neuron.

### Subset of stomatin domain genes expressed in mouse olfactory neurons

Previous work in *C. elegans* demonstrated a critical role for *mec-2* in mechanosensory behavior, and inspired follow-up work in mice demonstrating a role for *Stoml3* in mammalian mechanosensory behavior^11,12^. We similarly wished to determine whether the role for *mec-2* in olfaction represents a novel evolutionarily-conserved function for stomatin domain genes.

We first tested whether specific stomatin domain genes are selectively expressed in mouse olfactory receptor neurons (ORNs). To do so we analyzed RNA Seq data in which mouse olfactory receptor neurons were GFP labeled and FACS sorted, and compared this with similar data obtained from mouse motor neurons^9,28^ (Fig 3A-D). We confirmed that marker genes for the respective neuron classes are expressed in the expected cell types (Fig 3A, C), and found that stomatin domain genes exhibit distinct expression patterns (Fig 3B, D). We observed high expression of both *Stoml3* and *Stomatin* in ORNs, but not in motor neurons, while *Stoml1* and *Stoml2* are expressed in motor neurons but not olfactory neurons (Fig 3A-B). Because *Stoml3* displays the highest level of expression in ORNs, and because of a previous report demonstrating striking localization of *Stoml3* to ORN sensory cilia^8^, we prioritized *Stoml3* for further genetic and behavioral analysis.

**Figure 3:**
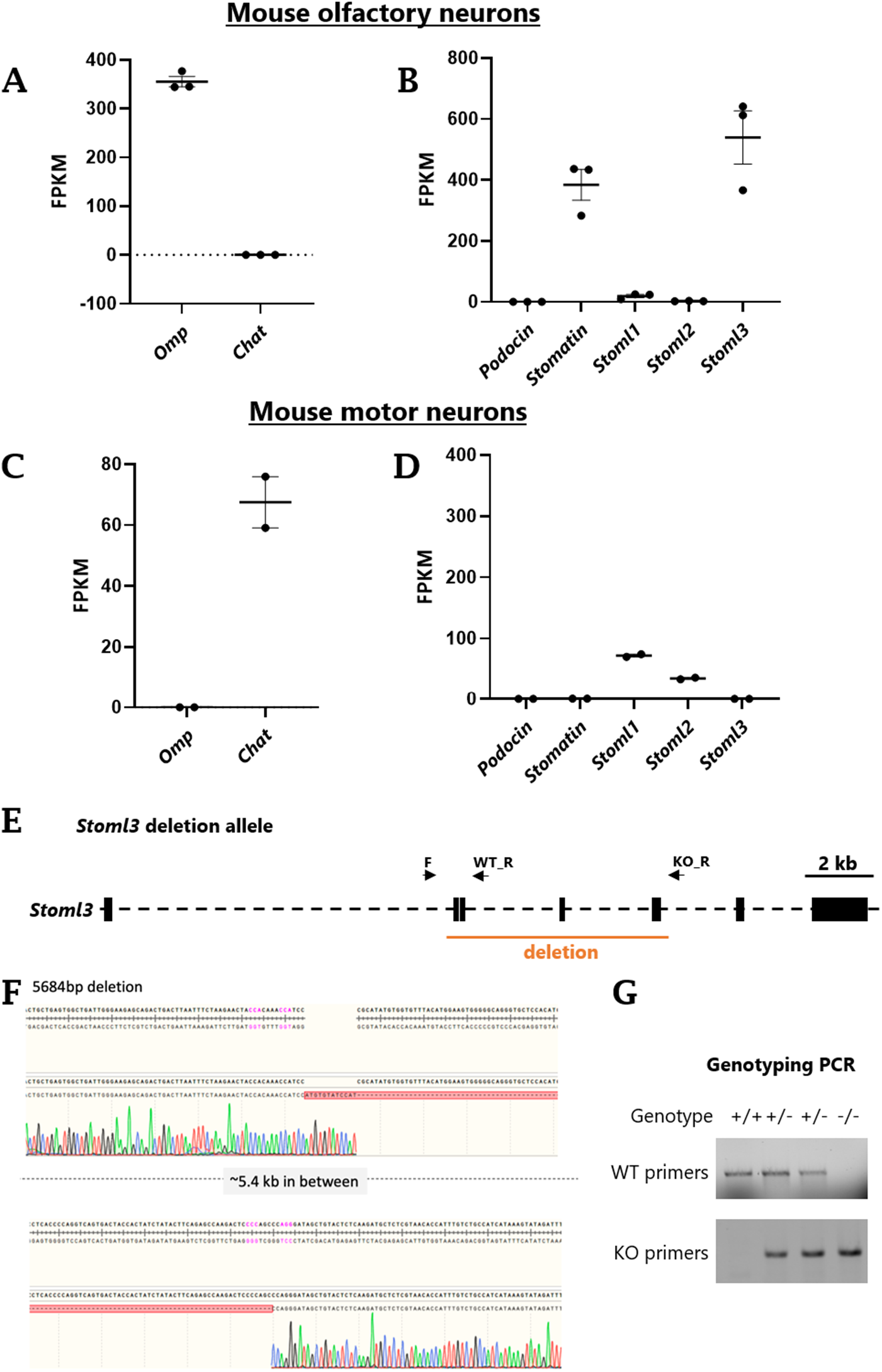
Stomatin domain genes expressed in mouse neurons, and *Stoml3* knockout mice. (A-D) Cell-specific sequencing of mouse olfactory neurons (A-B) and motor neurons (C-D). *Omp* is used as a positive marker for olfactory neurons, and *Chat* as a positive marker for cholinergic motor neurons. (E) *Stoml3* gene in mouse, indicating the region we deleted in orange, and the primer sets used to detect both the wild-type and mutant genomic DNA (black arrows). (F) Sanger sequencing confirming the ~5.4 kb deletion. (G) Genotyping gels for identifying homozygous *Stoml3* mutants and their wild-type littermates.

### *Stoml3* is required for olfactory behavior in mice

*Stoml3* KO mice have previously been generated^12^, but we were unable to obtain these mice, so we used CRISPR/Cas9 to generate a similar *Stoml3* KO mouse (Fig 3E-G). We removed the same two exons as in the published *Stoml3* KO, as well as two additional downstream exons. The deletion results in the loss of the conserved stomatin domain of *Stoml3* and introduction of downstream premature stop codons. As with the previous *Stoml3* KO mice, we found that *Stoml3*^-/-^ mice are born at Mendelian ratios and do not display overt developmental or behavioral defects.

To test whether *Stoml3* mutant mice are defective in olfactory behavior, we performed a number of standard olfactory behavioral assays on *Stoml3*^-/-^ mice and their wild-type littermates. We first tested the ability to detect and respond to olfactory stimuli that elicit innate behavioral responses in wild-type mice^29,30^. We measured time spent sniffing attractive odors compared to controls and found that both wild-type and *Stoml3* mutant mice spend more time sniffing attractive odors (Fig 4A-B), suggesting that *Stoml3* mutant mice retain the ability to detect odors. We next measured latency to find and eat a buried cereal pellet, because mice with impaired olfaction have been demonstrated to take longer to uncover buried food^31,32^. As with the innate olfactory assays, we found no differences between *Stoml3* and wild-type mice in latency to uncover the buried pellet (Figure 4C). Together these results suggest that loss of *Stoml3* does not affect innate olfactory response to attractive stimuli such as food odors.

**Figure 4:**
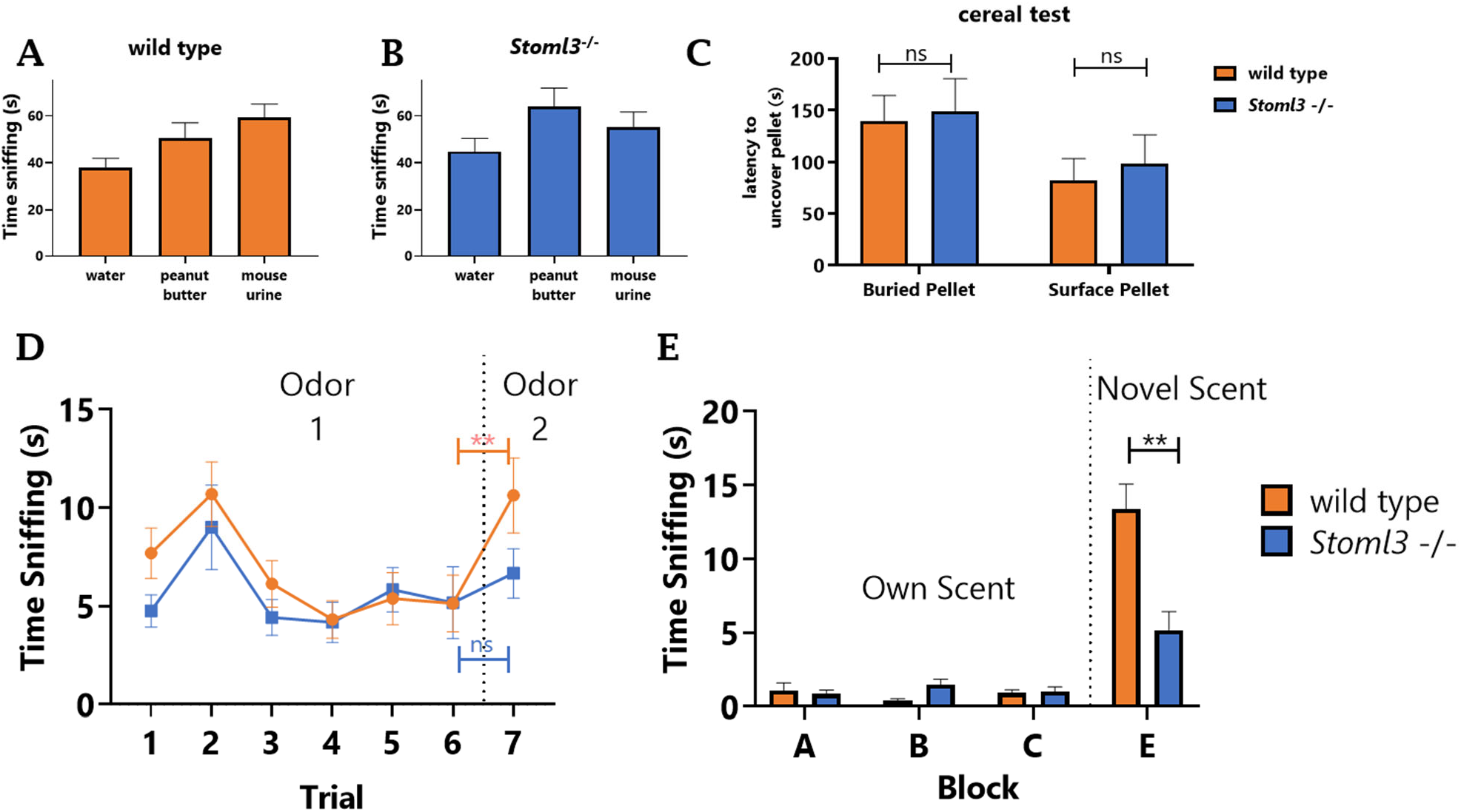
Olfactory assays for wild-type and *Stoml3* knockout mice. (A-B) Innate olfactory attraction tests. Mean time sniffing a scented block in wildtype mice (n = 16) and *Stoml3* knockout mice (n = 11) during the 3-min test period. Water was used as a control scent. Peanut butter and mouse urine were used as attractive scents. Both wild-type and *Stoml3* knock-out mice showed attraction response to peanut butter and mouse urine. (C) Buried cereal test. Mean time wild-type (n = 14) and *Stoml3* knock-out mice (n = 10) took to find the pellet. Surface pellet test is a positive control, confirming that the cereal pellet is attractive to the mice. ns represents no significant difference between wild types and *Stoml3* knockouts using Mann-Whitney U test. (D) Habituation/dishabituation test. Mean time sniffing a scented cartridge in wild type mice (n = 16) and *Stoml3* knockout mice (n = 12) during 30-sec test period across 7 trials. Almond extract was used on trial 1-6 and banana extract was introduced as a novel scent on trial 7. Significant difference of sniffing time in wild types is observed when novel scent is introduced on trial 7 (**represents p< 0.01), while no significant difference is observed between trial 6 and trial 7 in *Stoml3* knock-out mice. (E) Block test. Mean time sniffing home cage blocks A, B and C and novel block E in wildtype mice (n = 16) and *Stoml3* knockout mice (n = 12). Wild types spend more than 2-fold time exploring novel block (E) than *Stoml3* knockout mice (**represents p< 0.01, Mann-Whitney U test).

To test whether *Stoml3* knockout mice are able to distinguish between different odors, we performed a habituation/dishabituation test in which mice are habituated with one scent for six consecutive trials, then switched to a novel scent on the seventh trial^32^. We measured the time spent sniffing the odor cartridge during each trial. Both wild-type and *Stoml3* mutant mice display an initial interest in the odor, with sniffing time peaking during the second trial, followed by a period of habituation and decreased interest in the odor, as previously described^32^. On the seventh trial, the odor was changed to a novel scent. Wild-type mice spend more time sniffing the novel scent compared to a familiar scent (Fig 4D), as previously described^33^. In contrast, *Stoml3* mutant mice do not exhibit an increased interest in the novel scent in the seventh trial and spend the same amount of time investigating the novel scent in trial 7 as with the habituated scent in trial 6. This suggests that *Stoml3* mice, unlike wild-types, are unable to discriminate between familiar and novel scents (Fig. 4D).

Further evidence for the hypothesis that *Stoml3* mice are unable to discriminate between olfactory cues comes from experiments on social odors in the “block test.” In this test, we housed mice individually with several wooden blocks for 24 hours, allowing the blocks to acquire the odors of the home cage^32^. The next day, we removed and reintroduced these home cage blocks for six consecutive trials to measure investigation time of the familiar blocks. Mice became habituated to the presence of the blocks and spent minimal time investigating them by the sixth trial (Fig. S3). On the seventh trial, one of the blocks was swapped with a block from the cage of a different mouse, thus introducing a block with novel social odors. Wild-type mice spend a substantial amount of time sniffing of the novel-odor block, as previously described^33^ (Fig 4E). However, *Stoml3* mutants spend significantly less time sniffing the novel-odor block (Fig 4E), suggesting of a deficit in distinguishing between self- and non-self social odors.

Together, these experiments demonstrate that *Stoml3* knockout mice do not completely lose the ability to detect odors (Fig 4A-C). Rather, loss of *Stoml3* impairs the ability to discriminate between odors, including familiar versus novel odors (Fig 4D), and distinction between self and non-self odors (Fig 4E).

## DISCUSSION

Here we demonstrate that worms and mice express a small subset of stomatin domain genes in olfactory neurons. Of these, *mec-2* and *Stoml3* are required for proper olfactory behavior in both worms and mice. In worms, *mec-2* is required for chemotaxis, acts in a cell-autonomous manner, and multiple alternatively spliced *mec-2* isoforms can substitute for one another. In mice, *Stoml3* KO mice are able to detect odors, but unable to efficiently distinguish between odors. These experiments suggest that in addition to their well-established roles in mechanosensory behavior, *mec-2* and *Stoml3* have an additional shared role in olfactory behavior.

In a recent study, electrophysiological recordings from olfactory epithelium slices revealed that *Stoml3* KO ORNs exhibit modest reductions in spontaneous firing frequency^34^. Odor-evoked firing frequency is not affected, but spike duration and frequency are modestly reduced^34^. These electrophysiological properties of *Stoml3* KO OSNs could underly the deficits in olfactory behavior we describe here.

The precise biochemical role of stomatin domain proteins remains somewhat mysterious. A common theme is the modulation of ion channel function^35,36^, but the molecular mechanisms have yet to be fully elucidated. A model consistent with our data is that *mec-2* and *Stoml3* modulate ion channel activity in olfactory neurons in a manner required for their normal levels of activity. It will be interesting to test whether *mec-2/Stoml3* exhibit similar or distinct interactions with ion channels or other membrane proteins in mechanosensory versus olfactory neurons.

Finally, we now know that *mec-2* and *Stoml3* are required for at least two different sensory behaviors mediated by two different sensory neuron types (mechanosensory and olfactory). Are there additional neuronal roles yet to be uncovered? We note that in *C. elegans, mec-2* transcripts are expressed not only in the AWA and AWB olfactory neurons described here, but at comparably high levels in gustatory, thermosensory, and other specific neuronal types as well (Fig S3). It will be interesting to determine whether *mec-2* and/or *Stoml3* also play a role in these sensory activities as they do in mechanosensation and olfaction.

## ACKNOWLEDGEMENTS

Some strains were provided by the C.G.C., which is funded by NIH Office of Research Infrastructure Programs (P40 OD010440). ADN was supported by grants from the National Institute of Neurological Disorders and Stroke of the National Institutes of Health (R01NS111055) and the Welch Foundation (N-2042-20200401). Thank you to the entire Norris Lab, and to Megan Norris for helpful discussions, feedback, and critical reading/review of the manuscript.

## METHODS

### RNA Seq data

Raw fastq files were downloaded from the NCBI SRA for *C. elegans^21^* and mouse^9,28^ neuron-specific sequencing. Reads were mapped using STAR and alternative splicing mapped using JUM as previously reported^37–39^.

### Chemotaxis assays

Unseeded plates were divided into four quadrants, two with buffer and two with odorant, chemotaxis index = (# animals in two odorant quadrants)/(# animals in any of the four quadrants). The following modifications were made: animals were staged at larval L4 stage, M9 buffer was used for washes, animals were pelleted via gravity rather than centrifugation to minimize bacterial transfer, since *mec-2* loss-of-function mutants are lethargic in the presence of even trace amounts of bacterial food, and assays were extended to 2 hours to allow for a greater number of the uncoordinated worms to move out of the origin. *mec-2(e75)* is used as reference allele throughout.

### Smell on a stick

Worms were staged to larval L4s. 30+ worms of each genotype were transferred to unseeded plates. Each genotype was tested for lack of reaction to toothpick alone as well as toothpick and 70:30 95% EtOH: H2O (diluent). A 1:100,000 dilution of Octanol in 70:30 95% EtOH: H2O was made (testing solution). A sterile toothpick was dipped into the testing solution and placed just in front of a forward moving worm. The worm was scored for positive response (moving away from the test solution) or negative response (no response). Once a worm was tested it was removed to ensure it would not be re-assayed. n= 30 worms per genotype per assay, assays conducted in triplicate.

### Calcium Imaging

Calcium imaging was performed on young adult hermaphrodites on a microfluidic “worm chip” as previously described^26^, using the integrated transgene kyIs722 *str-2p*::GCaMP5(D380Y) to image GCaMP in the AWC(on) neuron. Isoamyl alcohol at 1:1,000 was used as olfactant.

### Mice

*Stoml3*^-/-^ mice were generated by deleting exon 2-5 of *Stoml3* using CRISPR/Cas9 genome editing. gRNAs used to generate deletion. Upstream: CACATATGCGGGATGGTTTG & TAAACACCACATATGCGGGA. Downstream: GAGCCAAGACTCCCCAGCCC & AGTACAGCTATCCCTGGGCT. Mice were housed in a temperature and light controlled room (12 h dark/light cycle) and all animal experiments were conducted in accordance with policies of NIH guide for the Care and Use of Laboratory Animals and Institutional Animal Care and Use Committee (IACUC) of Southern Methodist University. Adult mice (> 6-month-old) were used in this study, and both sexes of animals were used.

### Genotyping

DNA was extracted from tails of 19-21 days old mice. Wild-type and mutant mice were determined by PCR using primer set (Stoml3 common forward primer 5′ – TGTTCTCCCACATGCACACC – 3′, Stoml3 wild-type reverse primer 5′ – GGACCCTCATTAGATGCCCC – 3′, Stoml3 mutant reverse primer 5′ – GGCATCAGGTCCTCTGGAAC – 3′). Primer’s location on gene Stoml3 is illustrated in Figure 3E.

### Innate olfactory attraction test

Olfactory assays were conducted as previously described, with some modifications^29^. Mice were isolated and habituated with a block one day before the test day. On the test day, mice were transferred to a clean cage with a thin layer of bedding for at least 10 minutes for habituation. Bedding is essential in this step to help mice reduce the fear of open space. After habituation, a scented wood block with different test odor was introduced. Animals were recorded from the front side of the cage for 3 minutes. Sniffing time was defined as nasal contact with the block and was measured afterwards by analyzing the video. Odorants used were water (control, 80μL), peanut butter (10% w/v, 80 μL) and mouse urine (80 μL). Mouse urines were collected freshly from different litters and mixed well before use.

### Buried cereal test

Food restricted mice (90% of body weight) were used in buried cereal test to ensure mice were motivated to seek food. Mice were individually separated, and body weight was monitored every day before testing. A sweetened cereal was given to the tested mice before testing to overcome food neophobia. On all test days, mice were habituated for 1 hour in their cages without water or food. Clean cages were prepared with ~3 cm evenly distributed bedding, and 1 piece of sweetened cereal was buried 0.5 cm below the bedding. Mice were transferred from the habituation cage to the prepared new cage. A 5-minute timer was started when the mouse was introduced in the testing cage, and time was recorded when the mouse found the cereal and began eating it. If the mouse did not uncover the pellet within 5 min, 300 sec was recorded for that mouse. The buried cereal test was performed 5 days in a row, and the surface test was performed on the 6^th^ test (cereal was put on the surface of bedding). The time to uncover the buried pellet from day 3-5 was averaged and then compared between wild-type and mutant mice using a Mann-Whitney U test, similarly, time to uncover the surface pellet from 6^th^ day was compared between groups.

### Habituation/Dishabituation test

Mice were isolated in a clean cage overnight before testing. A tissue cartridge holding a non-scented cotton ball was placed in the cage to let mice get used to the novel item. On the testing day, mice were moved to the testing area without water and food for 1-hour habituation. After habituation, a scented tissue cartridge (noted as odor 1 in figure 4D) was placed into cage. Animals were recorded from the front side of the cage for 30 sec. This was repeated for 6 consecutive trials with odor 1, with inter-trial intervals of 5 minutes. On the 7^th^ trial, a novel odor (odor 2 in figure 4D) was introduced. Time sniffing was measured on each trial during the 30 sec test period. Almond extract (5 μL) was used as odor 1, banana extract (5 μL) was used as odor 2. Almond and banana scent were selected because they were considered as neutral odors for mice^40^.

### Block test

To measure ability to discriminate social odors, we performed the block test as previously described^32^. Each mouse was individually housed in a clean cage with bedding and 5 wood blocks labeled A-E at least 24 hours before testing. On the test day, all five blocks were removed from cage and placed into a sealed bag. Mice were transferred to testing area without food and water for 1-hour habituation. After habituation, blocks A-D from the same mouse were placed back into the cage. Mice were video recorded from front side of the cage for 30 sec. This procedure was repeated 6 times with at least 5 min interval. On the 7th trial, the same procedure was performed, but block D was replaced with block E from another mouse's cage so that A, B, C were home-cage blocks and block E was from other mouse's cage. Sniffing time of each block during the 30 sec test period from 7th trial was measured.

## SUPPLEMENTAL FIGURES

**Fig S1:**
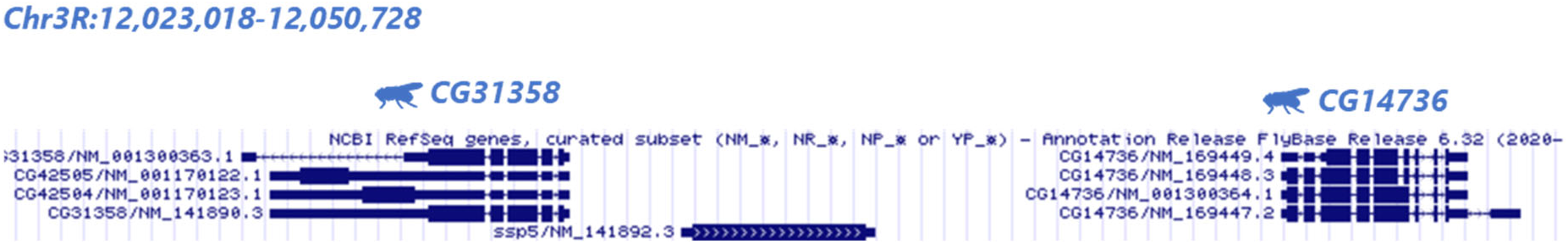
Additional cluster of *Drosophila Podocin* homologues on Chromosome 3.

**Fig S2:**
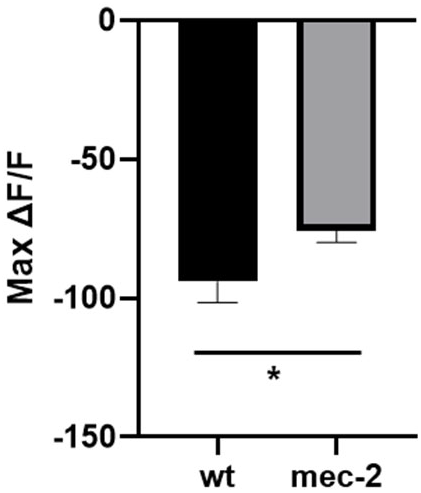
Reduced magnitude of calcium response in *mec-2(-)* mutants. Maximal ΔF/F GCaMP signals during odor presentation of Isoamyl alcohol (1:1,000) as displayed in Fig 2G. Unpaired two tail t-test, p<0.05.

**Figure S3:**
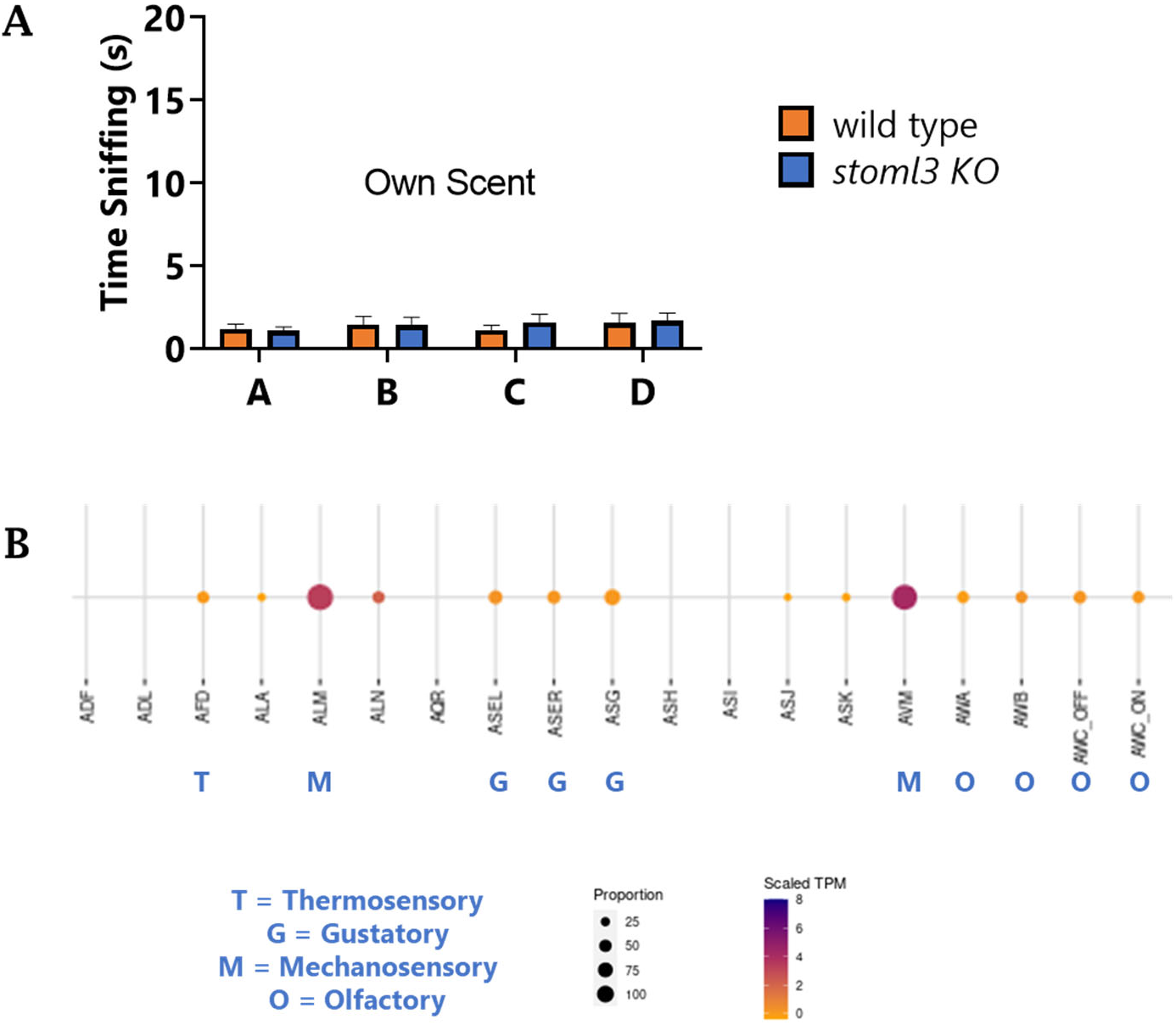
*Stoml3* behavior and *mec-2* expression. (A) In trial 6 of the block test, where no novel odor is present, time spent exploring the blocks is minimal. (B) Single-cell sequencing data from CenGEN consortium data on *mec-2* expression in representative neuron populations. Size of dot represents proportion of single cells in which *mec-2* was detected, and heatmap represents scaled TPM of the *mec-2* gene. A few sensory cell types are highlighted.

