## Supplemental Figures 1-3 for "A conserved role for stomatin domain genes in olfactory behavior"

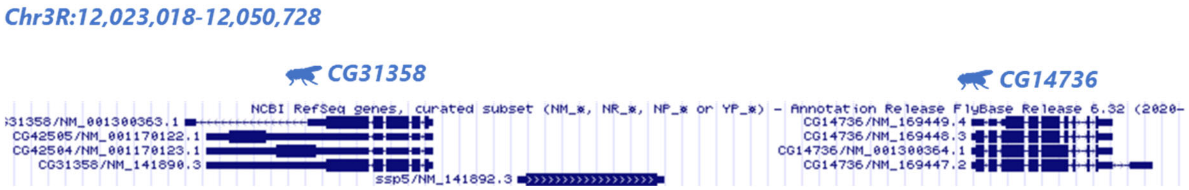


**Fig S1: Additional cluster of *Drosophila Podocin* homologues on Chromosome 3**

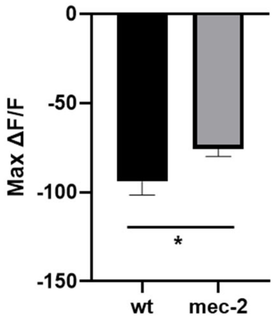


**Fig S2: Reduced magnitude of calcium response in *mec-2(-)* mutants.** Maximal ΔF/F GCaMP signals during odor presentation of Isoamyl alcohol (1:1,000) as displayed in Fig 2G. Unpaired two tail t-test, $p<0.05$.

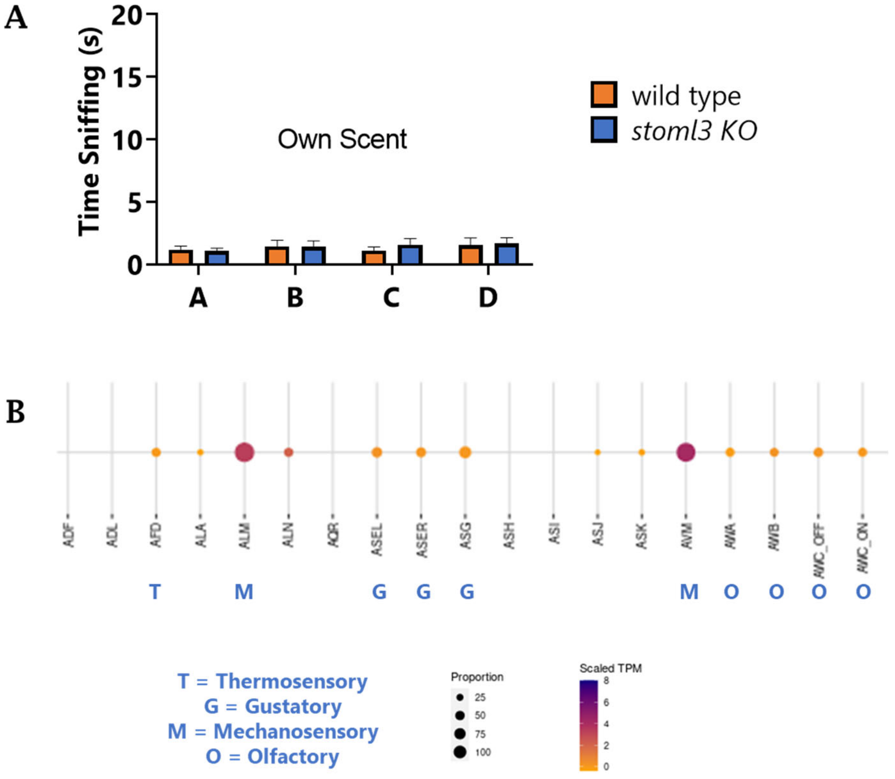


**Figure S3: *Stoml3* behavior and *mec-2* expression.** (A) In trial 6 of the block test, where no novel odor is present, time spent exploring the blocks is minimal. (B) Single-cell sequencing data from CenGEN consortium data on *mec-2* expression in representative neuron populations. Size of dot represents proportion of single cells in which *mec-2* was detected, and heatmap represents scaled TPM of the *mec-2* gene. A few sensory cell types are highlighted.
